# EEG resting-state large-scale brain network dynamics are related to depressive symptoms

**DOI:** 10.1101/619031

**Authors:** Alena Damborská, Miralena I. Tomescu, Eliška Honzírková, Richard Barteček, Jana Hořínková, Sylvie Fedorová, Šimon Ondruš, Christoph M. Michel

## Abstract

**Background:** The few previous studies on resting-state EEG microstates in depressive patients suggest altered temporal characteristics of microstates compared to those of healthy subjects. We tested whether resting-state microstate temporal characteristics could capture large-scale brain network dynamic activity relevant to depressive symptomatology.

**Methods:** To evaluate a possible relationship between the resting-state large-scale brain network dynamics and depressive symptoms, we performed EEG microstate analysis in patients with moderate to severe depression within bipolar affective disorder, depressive episode, and periodic depressive disorder, and in healthy controls.

**Results:** Microstate analysis revealed six classes of microstates (A-F) in global clustering across all subjects. There were no between-group differences in the temporal characteristics of microstates. In the patient group, higher symptomatology on the Montgomery-Åsberg Depression Rating Scale, a questionnaire validated as measuring severity of depressive episodes in patients with mood disorders, correlated with higher occurrence of microstate A (Spearman’s rank correlation, r = 0.70, p < 0.01).

**Conclusion:** Our results suggest that the observed interindividual differences in resting-state EEG microstate parameters could reflect altered large-scale brain network dynamics relevant to depressive symptomatology during depressive episodes. These findings suggest the utility of the microstate analysis approach in an objective depression assessment.

## 1 Introduction

Major depressive disorder (MDD) and bipolar disorder are among the most serious psychiatric disorders with high prevalence and illness-related disability (Andrade et al., 2003; Eaton et al. 2012; Cloutier et al., 2018). Despite growing evidence for the spectrum concept of mood disorders (Angst et al., 2018), and even with the advanced neuroimaging methods developed in recent years, the underlying pathophysiological mechanisms of depression remain poorly understood. Evidence across resting-state functional magnetic resonance (fMRI) studies consistently points to an impairment of large-scale resting-state brain networks in MDD rather than a disruption of discrete brain regions (Gong and He, 2015; Iwabuchi et al., 2015; Kaiser et al.,2015; Peng et al.,2015). Consistent with the neurobiological model of depression (Mayberg et al., 1997), numerous resting-state fMRI studies show decreased frontal cortex function and increased limbic system function in patients with MDD (Fischer et al., 2016). Functional abnormalities in large-scale brain networks include hypoconnectivity within the frontoparietal network (Kaiser et al., 2015) and the reward circuitry, centered around the ventral striatum (Satterthwaite et al., 2015). Reduced functional connectivity in first-episode drug-naïve patients with MDD was also recently reported between the frontoparietal and cingulo-opercular networks (Wu et al., 2016). Moreover, hyperconnectivity of the default mode network (Greicius et al., 2007) and amygdala hyperconnectivity with the affective salience network (Price and Drevets, 2010; Hamilton et al., 2013) were shown to be characteristic features of depression.

In general, large-scale networks dynamically re-organize themselves on sub-second temporal scales to enable efficient functioning (de Pasquale et al., 2017; Bressler and Menon, 2010). Fast temporal dynamics of large-scale neural networks, not accessible with the low temporal resolution of the fMRI technique, can be investigated by analyzing the temporal characteristics of “EEG microstates” (Van de Ville et al., 2010; Michel and Koenig, 2018). Scalp EEG measures the electric potential generated by the neuronal activity in the brain with a temporal resolution in the millisecond range. A sufficient number of electrodes distributed over the scalp, i.e. high density-EEG (HD-EEG), allows for the reconstruction of a scalp potential map representing the global brain activity (Michel and Murray, 2012). Any change in the map topography reflects a change in the distribution and/or orientation of the active sources in the brain (Lehmann, 1987). Already in 1987, Lehmann et al. observed that in spontaneous resting-state EEG, the topography of the scalp potential map remains stable for a short period of time and then rapidly switches to a new topography in which it remains stable again. Ignoring map polarity, the duration of these stable topographies is around 80-120 ms. Lehmann called these short periods of stability EEG microstates and attributed them to periods of synchronized activity within large-scale brain networks. For a recent review, see (Michel and Koenig, 2018). Assessment of the temporal characteristics of these microstates provides information about the dynamics of large-scale brain networks, because this technique simultaneously considers signals recorded from all areas of the cortex. Since the temporal variation in resting-state brain network dynamics may be a significant biomarker of illness and therapeutic outcome (Hutchison et al., 2013; Chang and Glover, 2010; Honey et al., 2007), microstate analysis is a highly suitable tool for this purpose.

Numerous studies demonstrated changes in EEG microstates in patients with neuropsychiatric disorders such as schizophrenia, dementia, panic disorder, multiple sclerosis, and others (for reviews see Khanna et al., 2015; Michel and Koenig, 2018). Despite the potential of microstate analysis for detecting global brain dynamic impairment, microstates were not investigated in depressive patients, except for three studies that provided inconsistent results. Using adaptive segmentation of resting state EEG in depressive patients, two early studies showed abnormal microstate topographies and reduced overall average microstate duration (Strik et al., 1995) but unchanged numbers of different microstates per second (Ihl and Brinkmeyer, 1999). In a more recent study using a topographical atomize-agglomerate hierarchical clustering algorithm, abnormally increased overall microstate duration and decreased overall microstate occurrence per second were reported in treatment-resistant depression (Atluri et al., 2018).

A better understanding of disruption and changes in brain network dynamics in depression is critical for developing novel and targeted treatments, e.g. deep brain stimulation in treatment-resistant depression (Drobisz and Damborská 2019). Furthermore, microstate features reflecting the disruption of brain network dynamics might be later tested as candidate biomarkers of depressive disorder and predictors of treatment response. Thus, the main goal of our study was to explore how resting-state microstate dynamics are affected in depressive patients as compared to healthy individuals. We hypothesized that patients with depression will show different microstate dynamics than healthy controls in terms of the temporal characteristics of EEG microstates such as duration, coverage, and occurrence. We also hypothesized that microstate dynamics will be related to the overall clinical severity of depression.

## 2 Materials and Methods

### 2.1 Subjects

Data was collected from 19 depressive patients (Age in years: mean = 53.0, standard deviation = 9.8; 6 females) and 19 healthy control (HC) subjects (Age in years: mean = 51.4, standard deviation = 9.1; 6 females). There were no differences in gender and an independent sample *t*-test (*t*-value (df 36) = 0.45, p > 0.05) also showed no significant difference in age between the two groups. The patients were recruited by the Department of Psychiatry at the University Hospital Brno based on the clinical evaluation by two board-certified psychiatrists and according to the Diagnostic and Statistical Manual (DSM-V) criteria using the Mini International Neuropsychiatric interview (M.I.N.I.). All patients were examined during their first week of hospitalization. All patients met the criteria for at least a moderate degree of depression within the following affective disorders: bipolar affective disorder (F31), depressive episode (F32), and periodic depressive disorder (F33). Exclusion criteria for patients were any psychiatric or neurological comorbidity, IQ < 70, organic disorder with influence on the brain function, alcohol dependence or other substance dependence. All patients were in the on-medication state. The patient characteristics are summarized in Table 1. Control subjects were recruited by general practitioners from their database of clients. Control subjects underwent the M.I.N.I. by board-certified psychiatrists, to ensure that they had no previous or current psychiatric disorder according to the DSM-V criteria. The scores on the Montgomery-Åsberg Depression Rating Scale (MADRS) (Williams and Kobak, 2008) and the Clinical Global Impression (CGI) (Guy, 1976) tests were measured in patients to assess the severity of the depressive episode. The status of depression was further described with lifetime count of depressive episodes. Medication in 24 hours preceding the EEG examination was also recorded (see Table 1). This study was carried out in accordance with the recommendations of Ethics Committee of University Hospital Brno with written informed consent from all subjects. All subjects gave written informed consent in accordance with the Declaration of Helsinki. The protocol was approved by the Ethics Committee of University Hospital Brno, Czech Republic.

**Table 1.** Patient characteristics

| Patient | Diagnose | Number of episodes | MADRS score | CGI score | BZD | AD/AP/MS |
| --- | --- | --- | --- | --- | --- | --- |
| 1 | F31 | 3 | 27 | 4 | 2 | 3 |
| 2 | F32 | 1 | 24 | 5 | 0 | 2 |
| 3 | F32 | 1 | 15 | 4 | 2 | 2 |
| 4 | F31 | 5 | 39 | 6 | 0 | 2 |
| 5 | F33 | 3 | 18 | 4 | 0 | 1 |
| 6 | F33 | 2 | 9 | 3 | 1.33 | 1 |
| 7 | F32 | 1 | 24 | 4 | 1.33 | 3 |
| 8 | F31 | 4 | 29 | 5 | 2 | 2 |
| 9 | F33 | 2 | 36 | 6 | 1 | 4 |
| 10 | F33 | 3 | 21 | 4 | 1 | 1 |
| 11 | F33 | 2 | 38 | 5 | 6 | 4 |
| 12 | F33 | 2 | 39 | 5 | 3 | 4 |
| 13 | F32 | 1 | 21 | 5 | 2 | 4 |
| 14 | F33 | 5 | 32 | 5 | 0 | 3 |
| 15 | F33 | 2 | 38 | 6 | 3 | 4 |
| 16 | F32 | 1 | 37 | 6 | 2 | 4 |
| 17 | F33 | 3 | 18 | 4 | 0 | 4 |
| 18 | F31 | 2 | 28 | 4 | 0 | 4 |
| 19 | F31 | 11 | 23 | 4 | 1 | 4 |

### 2.2 EEG recording and pre-processing

Subjects were sitting in a comfortable upright position in an electrically shielded room with dimmed light. They were instructed to stay as calm as possible, to keep their eyes closed and to relax for 15 minutes. They were asked to stay awake. All participants were monitored by the cameras and in the event of signs of nodding off or EEG signs of drowsiness detected by online visual inspection, the recording was stopped. The EEG was recorded with a high density 128-channel system (EGI System 400; Electrical Geodesic Inc., OR, USA), a sampling rate of 1kHz, and Cz as acquisition reference. Five minutes of the EEG data were selected based on visual assessment of the artifacts. The EEG was band-pass filtered between 1 and 40 Hz. Subsequently, in order to remove ballistocardiogram and oculo-motor artifacts, infomax-based Independent Component Analysis (Jung et al., 2000) was applied to all but one or two channels rejected due to abundant artifacts. Only components related to ballistocardiogram, saccadic eye movements, and eye blinking were removed based on the waveform, topography, and time course of the component. The cleaned EEG recording was down-sampled to 125 Hz and the previously identified noisy channels were interpolated using a three-dimensional spherical spline (Perrin et al., 1989), and re-referenced to the average reference. For subsequent analyses, the EEG data was reduced to 110 channels to remove muscular artifacts originating in the neck and face. All the preprocessing steps were done using the freely available Cartool Software 3.70, programmed by Denis Brunet Cartool (https://sites.google.com/site/cartoolcommunity/home) and MATLAB.

### 2.3 Microstate analysis

The microstate analysis (see Fig. 1) followed the standard procedure using *k*-means clustering method to estimate the optimal set of topographies explaining the EEG signal (Brunet et al., 2011; Murray et al, 2008; Pascual-Marqui et al., 1995). The polarity of the maps was ignored in this clustering procedure. To determine the optimal number of clusters, we applied a meta-criterion that is a combination of seven independent optimization criteria (for details see Brechet et al., in press). In order to improve the signal-to-noise ratio, only the data at the time points of the local maximum of the Global Field Power (GFP) were clustered (Pascual-Marqui et al., 1995; Koenig et al., 2002; Britz et al, 2010, Tomescu et al., 2014). The GFP is a scalar measure of the strength of the scalp potential field and is calculated as the standard deviation of all electrodes at a given time point (Michel et al., 1993; Brunett et al., 2011; Murray et al., 2008). The cluster analysis was first computed at the individual level and then at global level across all participants (patients and controls).

**Figure 1.**
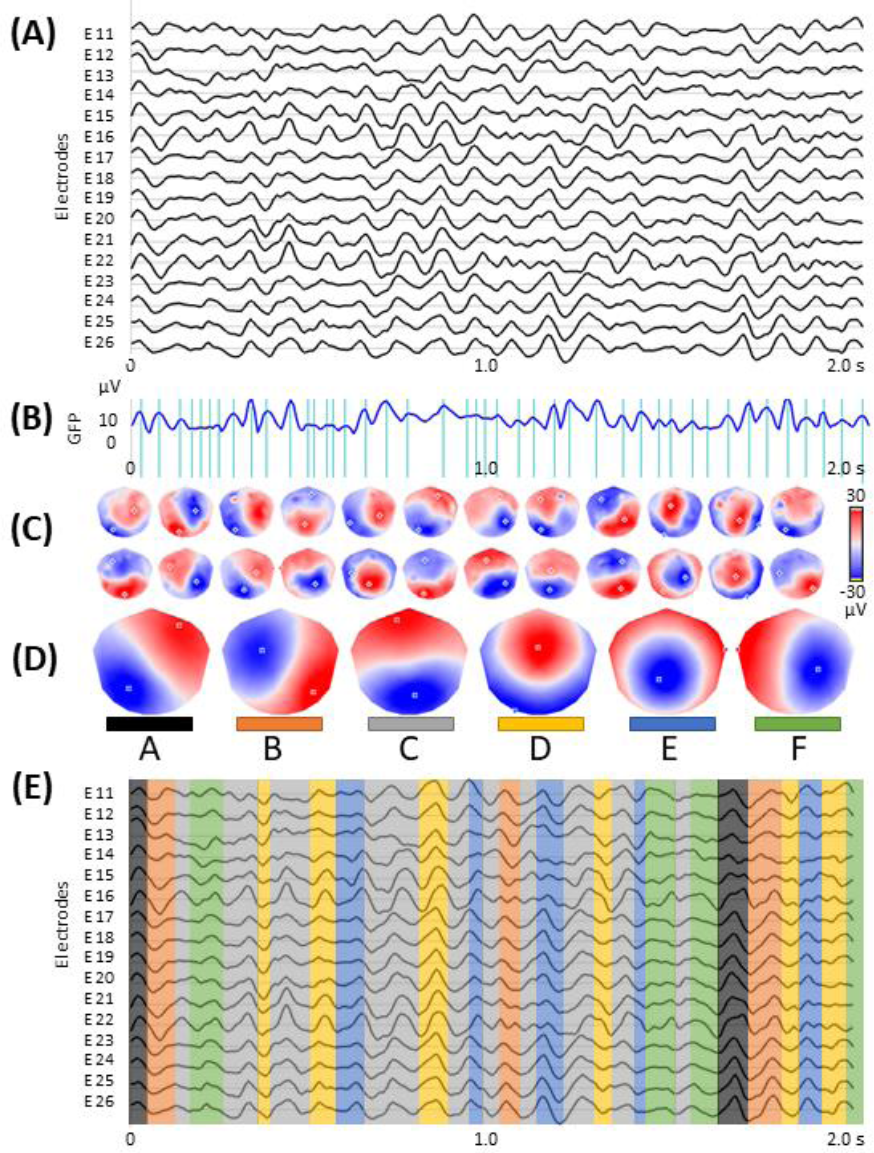
Microstate analysis: **(A)** resting-state EEG from subsample of 16 out of 110 electrodes; **(B)** global field power (GFP) curve with the GFP peaks (vertical lines) in the same EEG period as shown in (A); **(C)** potential maps at successive GFP peaks, indicated in (B), from the first 1 s period of the recording; **(D)** set of six cluster maps best explaining the data as revealed by K-means clustering of the maps at the GFP peaks; **(E)** the original EEG recording shown in (A) with superimposed color-coded microstate segments. Note that each time point of the EEG recording was labelled with the cluster map, shown in (D), with which the instant map correlated best. The duration of segments, occurrence, and coverage for all microstates were computed on thus labeled EEG recording.

Spatial correlation was calculated between every map identified at the global level and the individual subject’s topographical map in every instant of the pre-processed EEG recording. Each continuous time point of the subject’s EEG (not only the GFP peaks) was then assigned to the microstate class of the highest correlation, again ignoring polarity (Brunet et al., 2011; Brechet et al., in press; Michel and Koenig, 2018; Santarnecchi et al., 2017). Temporal smoothing parameters (window half size = 3, strength (Besag Factor) = 10) ensured that the noise during low GFP did not artificially interrupt the temporal segments of stable topography (Brunet et al., 2011; Pascual-Marqui et al., 1995). For each subject, three temporal parameters were then calculated for each of the previously identified microstates: (i) occurrence, (ii) coverage, and (iii) duration. Occurrence indicates how many times a microstate class recurs in one second. The coverage represents the summed amount of time spent in a given microstate class. The duration in milliseconds for a given microstate class indicates the amount of time that a given microstate class is continuously present. In order to assess the extent to which the representative microstate topographies explain the original EEG data, the global explained variance (GEV) was calculated as the sum of the explained variances of each microstate weighted by the GFP. Microstate analysis was performed using the freely available Cartool Software 3.70, programmed by Denis Brunet Cartool (https://sites.google.com/site/cartoolcommunity/home).

### 2.4 Statistical analysis

To investigate group differences, independent *t*-tests were used for temporal parameters of each microstate. Comparisons were corrected using the false discovery rate (FDR) method (Benjamini, 2010). In order to evaluate the possible relation of microstate dynamics to severity of depression, we computed Spearman’s rank correlation coefficients of all microstate parameters with the MADRS and CGI scores and number of episodes. In order to evaluate possible influence of medication on microstate dynamics, we calculated Spearman’s rank correlation coefficients between all microstate parameters and medication, that patients received during 24 hours preceding the EEG measurement. Intake of antidepressants, antipsychotics and mood stabilizers was indicated as a single ordinal variable taking into account the number of medicaments and their dosages. Intake of benzodiazepines was expressed with the benzodiazepine equivalent dose (Bazire, 2014). A significance level of α < 0.01 was used for all correlations. Statistical evaluation of the results was performed by the routines included in the program package Statistica’13 (1984-2018, TIBCO, Software Inc, Version 13.4.0.14).

## 3 Results

The meta-criterion used to determine the most dominant topographies revealed six microstates explaining 82.6 % of the global variance. Four topographies resembled those previously reported in the literature as A, B, C, and D maps (Michel and Koenig, 2018; Koenig et al., 2002; Britz 2010; Atluri et al., 2018) and two topographies resembled the recently identified (Custo et al., 2017) resting-state microstate maps. We labeled these maps as A – D, in accordance with previous literature, and as E and F (Fig. 2).

**Figure 2.**
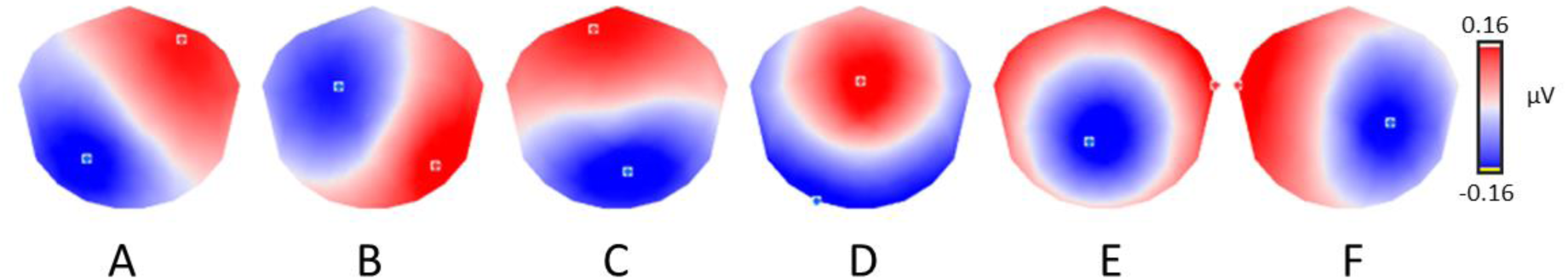
The six microstate topographies identified in the global clustering across all subjects.

The groups did not differ in any temporal parameter in any microstate. The depressive group was indistinguishable from the control group (all absolute *t*-values < 2.5). The FDR-corrected p-values (six comparisons for the six microstate classes) were not significant between the patients and controls for any microstate in the duration (A: p=0.39; B: p=0.39; C: p= 0.30; D: p=0.39; E: p=0.77; F: p=0.68), occurrence (A: p=0.13; B: p=0.92; C: p= 0.92; D: p=0.92; E: p=0.13; F: p=0.29), or coverage (A: p=0.44; B: p=0.75; C: p= 0.44; D: p=0.75; E: p=0.16; F: p=0.44).

The results of Spearman’s rank correlation revealed a positive association of the depression severity with the presence of microstate A but not with the presence of other microstates. The occurrence of microstate A significantly correlated with the MADRS scores (r = 0.70, p < 0.01; Figure 3), but not with the CGI score (r = 0.40) or the number of episodes (r = 0.08). There were no significant associations between the depression severity and the duration or coverage of microstate A (all absolute r-values < 0.55).

**Figure 3.**
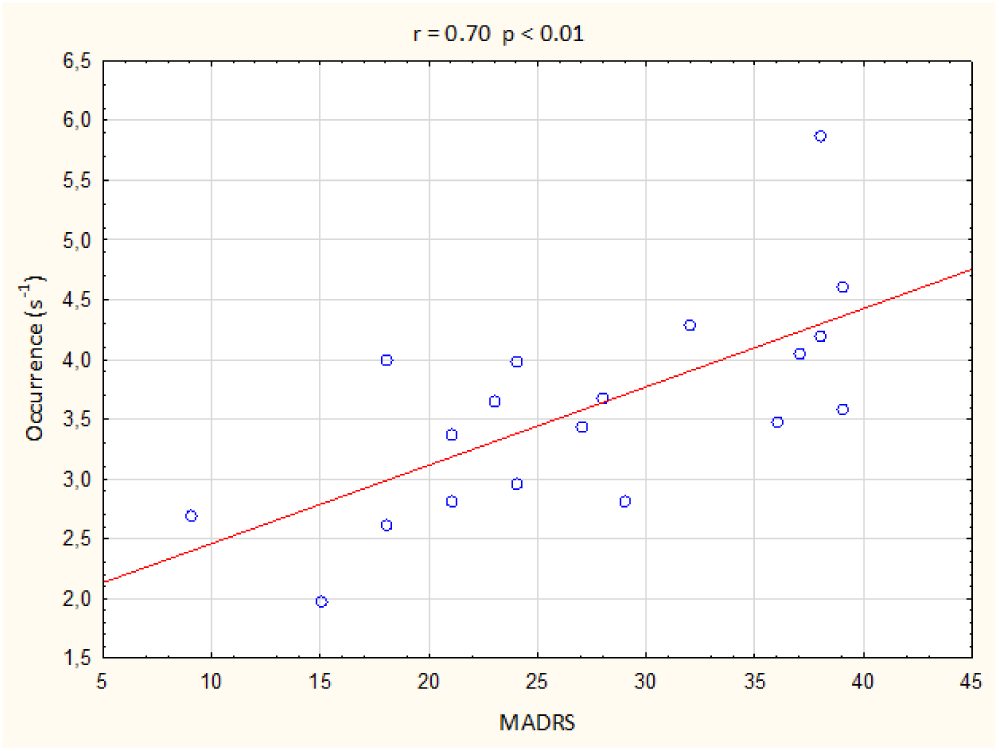
Correlations between the occurrence of microstate A and Montgomery-Åsberg Depression Rating Scale (MADRS) score.

The results of Spearman’s rank correlation revealed a significant positive association between the medication status and the presence of microstate E but not with the presence of other microstates. The occurrence of microstate E significantly correlated with the intake of antidepressants, antipsychotics, and mood stabilizers (r = 0.65, p < 0.01; Figure 4), but not with the intake of benzodiazepines (r = 0.20). There were no significant associations between the medication status and the duration or coverage of microstate E (all absolute r-values < 0.45).

**Figure 4.**
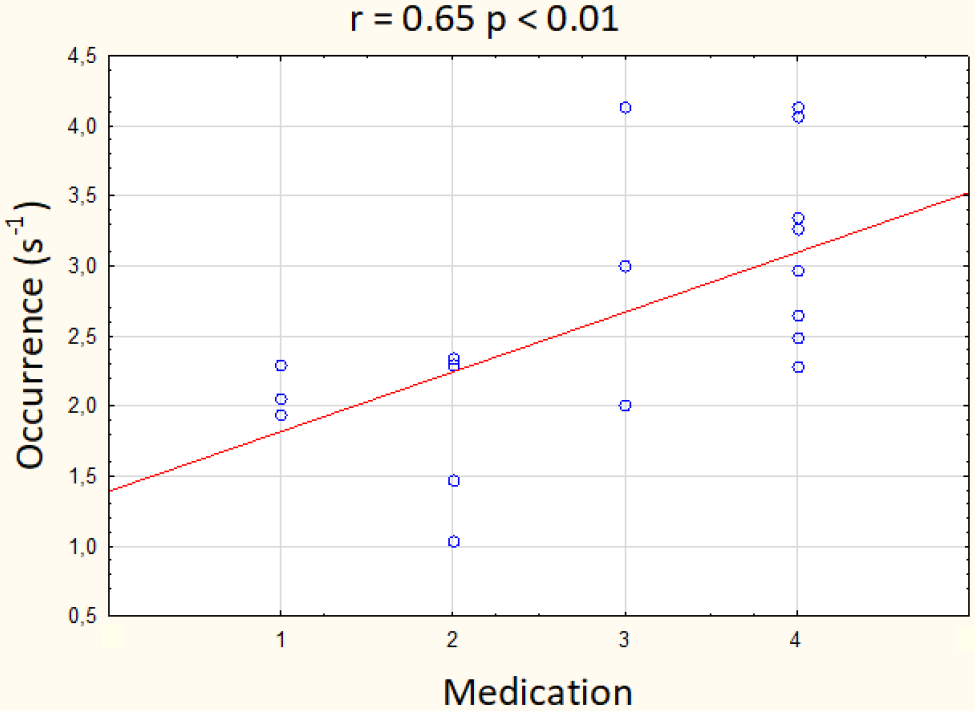
Correlations between the occurrence of microstate E and the intake of antidepressants, antipsychotics, and mood stabilizers. Medication scale: 1 – one medication in sub-therapeutic doses, 2 – one medication in therapeutic doses, 3 – combination of medications with one in therapeutic doses, 4 – combination of medications with more than one in therapeutic doses

## 4 Discussion

In this report, the dynamics of resting-state large-scale brain network activity are depicted in the form of functional EEG brain microstates. We demonstrated that microstate temporal dynamics are sensitive to interindividual differences in depressive symptom severity in patients with moderate to severe depression. Particularly, we showed that severity of depressive symptoms correlated with higher occurrence of the microstate A. This finding suggests that microstate analysis-based neural markers might represent a largely untapped resource for understanding the neurobiology of depression. The present study is the first in a planned longitudinal study series with depressive patients recruited at the University Hospital Brno that will help further investigate the microstate parameters as possible predictors of treatment response to both medication and neurostimulation methods including the electroconvulsive therapy.

Only three studies with inconsistent results examined microstate duration and occurrence in depressive patients. While the earlier studies showed reduced duration (Strik et al., 1995) and unchanged occurrence (Ihl and Brinkmeyer, 1999) in depression, the more recent study reported longer duration and lower occurrence of microstates in treatment-resistant depressed patients than in healthy subjects (Atluri et al., 2018). The authors suggested that the decreased occurrence and increased duration in microstates could be linked to the neurotropic medications previously taken by patients resistant to antidepressant treatment. Namely in the cited study, the between group differences were greater following the seizure therapy. The authors hypothesized that the antidepressant exposure may have an effect on global brain dynamics even in pharmaco-resistant patients. They further argue that antidepressants and seizure therapy may modulate global brain dynamics in a similar manner, i.e. decrease the occurrence and increase the duration of microstates.

In the current study, we found an increased microstate A occurrence with depression only as an effect related to the symptom severity and not as a group difference. This finding is, however, consistent with the previously reported lowering of microstate occurrence following seizure therapy that in fact might represent a normalization of occurrence with successful treatment (Atluri et al., 2018).

Due to fundamental differences in methodological approaches, it is difficult to compare our findings with the three previous microstate studies on depression. The discrepancies among methodological approaches employed include different frequency band examined, different clustering algorithms applied, different numbers of maps used for backfitting to the EEG, and analyzing all data points (e.g. in current study) or only those with the local maxima of the global field power (e.g. in Alturi et al., 2018). Discrepant findings may also reflect the pathophysiologic heterogeneity of depression. Similarly to the current sample, the experimental group in the study by Strik et al. (1995) included depressive patients who met the criteria for unipolar or bipolar mood disorders or for dysthymia. The other two studies both focused on unipolar depression (Ihl and Brinkmeyer, 1999; Atluri et al., 2018), the more recent one was even restricted only to the treatment-resistant form of depression (Atluri et al., 2018). With respect to the symptom variations in patients meeting criteria for depression, currently based solely on the clinical interviews and diagnostic questionnaires, such heterogeneity in findings could be expected.

In the current study, the topography of microstate A strongly resembled the topography of one of the four canonical microstates, i.e. microstate A, earlier described in the literature (Khanna et al., 2015; Michel and Koenig, 2018). Using resting-state fMRI, this microstate was previously linked to the auditory brain network (Britz et al., 2010), involving bilateral superior and middle temporal gyri, regions associated with phonological processing (Buchsbaum et al., 2001). In addition to this indirect identification of involved brain structures, the sources generating microstate scalp topographies were directly estimated (Custo et al., 2017; Brechet et al., in press). The left temporal lobe and left insula were identified as the major generators of microstate A (Custo et al., 2017). Additionally, left-lateralized activity in the medial prefrontal cortex and the occipital gyri was most recently reported to underlie this microstate (Brechet et al., in press).

Evidence from the meta-analysis of functional neuroimaging studies suggests resting-state functional alterations in first-episode drug-naïve MDD patients in the fronto-limbic system, including the dorsolateral prefrontal cortex and putamen, and in the default mode network, namely the precuneus and superior and middle temporal gyri (Zhong et al., 2016). Altered activity in the superior temporal gyrus in patients with MDD was reported repeatedly in fMRI studies (Ke et al., 2016; Liu et al., 2010; Shen et al.,2014; Tadayonnejad et al., 2015) and was also suggested to be responsible for the abnormal processing of negative mood and cognition in first-episode, drug-naïve patients with MDD (Zhong et al., 2016). Our findings of positive associations of depressive symptoms with the occurrence of microstate A that is related to temporal lobe activity are thus in line with these studies.

It has been shown that benzodiazepines and antipsychotics may modulate microstate dynamics (Kinoshita et al., 1995). Accordingly, we observed the effect of medication on the presence of microstate E. The topography of this microstate strongly resembled one of the newly reported microstates, the generators of which were identified in the dorsal anterior cingulate cortex, superior and middle frontal gyri, and insula (Custo et al., 2017). The cingulo-opercular network (CON), comprising regions in the thalamus as well as frontal operculum/anterior insula and anterior cingulate cortex, is considered to have a central role in sustaining alertness (Coste and Kleinschmidt, 2016) or in general for maintaining perceptual readiness (Sadaghiani and D’Esposito, 2015). An important role in the pathophysiological mechanisms of depression was suggested for the CON, whose disrupted functional connectivity was observed in first-episode drug-naïve patients with MDD (Wu et al., 2016). Our findings of association between the occurrence of microstate E and on-medication status might be related to the pharmacological effect on the activity of structures constituting the CON.

In the current study, we decided to use the resting-state condition rather than employing any cognitive task. Depression affects not only emotional and cognitive mental operations but also motivational processes. Therefore, the task performance differences between patients and healthy controls may relate to different levels of motivation rather than information processing per se. Using a resting-state condition makes it possible to avoid some task-related confounds and makes the application of non-invasive neuroimaging techniques a powerful tool for measureing baseline brain activity (Gusnard and Raichle, 2001). Moreover, if the research outputs such as those presented here lead to developing a new diagnostic tool for depressive disorder, such a tool, based on evaluating the resting-state scalp EEG, will be easy to use and require only minimal cooperation from the patients.

It is important to note that our data may have limitations. First, our sample included mixed diagnoses, with both bipolar and unipolar disorders. The observed relationship between the microstate A occurrence and depressive symptomatology should therefore be considered as a state rather than as a trait marker of depression. Second, the low sample size and great variability in medication made it impossible to examine any potential influence of medication on the microstate parameters by comparing patients receiving a specific drug with those not receiving it. To summarize the various medications, an ordinal variable was used that is only a rough measurement of medication usage. Therefore, the observed relationship between the microstate E occurrence and medications should be viewed with caution.

## 5 Conclusions

The study presented here provides insights into global brain dynamics of the resting-state in depressive patients. The identified depressive symptom related changes in resting-state large-scale brain dynamics suggest the utility of the microstate analysis approach in an objective depression assessment. To test the observed microstate changes as possible biomarkers of illness and/or treatment response at individual level is the next step for future research in depressive patients.

## 8 Conflict of Interest

The authors declare that the research was conducted in the absence of any commercial or financial relationships that could be construed as a potential conflict of interest.

## 9 Author Contributions

AD – designed the study, performed the analysis, and wrote the initial draft; MT – served as consultant for the data analysis; RB and JH – were responsible for patient recruitment and clinical assessment; EH – collected the HD-EEG data; SF and ŠO – were involved in the clinical assessment; CM – served as an advisor and was responsible for the overall oversight of the study. All authors revised the manuscript.

## 10 Funding

This project received funding from the European Union Horizon 2020 research and innovation program under the Marie Skłodowska-Curie grant agreement No. 739939. The study was also supported by Ministry of Health, Czech Republic - conceptual development of research organization (University Hospital Brno - FNBr, 65269705). These funding sources had no role in the design, collection, analysis, or interpretation of the study.

## 11 Acknowledgments

The authors wish to thank Anne Meredith Johnson for providing language help.

## 13 Data Availability Statement

The raw data supporting the conclusions of this manuscript will be made available by the authors, without undue reservation, to any qualified researcher.

